# Benchmarking sample mispairing detection tools in complex cancer datasets

**DOI:** 10.64898/2026.11.23.768692

**Authors:** Emma Lathouwers, Saloni Dixon, Grace Y. Akatsu, Ariel A. Hippen, Laurie Grieshober, Euihye Jung, Lukas M. Weber, Ronny Drapkin, Stephanie C. Hicks, Jennifer A. Doherty, Casey S. Greene

## Abstract

Sample mispairs are common and potentially costly mistakes, and therefore checking for sample mispairing is a crucial quality-control step in multimodal genomic studies. This study presents an independent benchmark of sample mispairing detection tools in multimodal transcriptomic cancer datasets. In pseudobulk experiments with known sample pairings, four tools achieved near-perfect F1 scores across datasets. In real-data experiments, three tools performed particularly well, and sample mispairs were identified in three of five datasets. Finally, a high-grade serous ovarian carcinoma dataset is reconstructed: resolving mispairs between two bulk RNA-sequencing modalities, resolving mispairs between pooled single-cell and bulk RNA-seq data, and reestablishing the connection between the transcriptomic data and patient metadata using genetic ancestry inference.

**Author summary:** Sample mislabeling errors are common and potentially costly mistakes in sequencing studies. With one study finding 40% of transcriptomic datasets contain mislabeling errors and other studies placing a lower bound on the per-sample error rate around 1% These errors can reduce the ability to detect biologically meaningful signals, contribute to incorrect conclusions, and waste experimental and computational resources. In multimodal sequencing studies, mislabeling can lead to sample mispairs: Samples are attributed to the same patient when they do not originate from the same patient. We tested available sample mispairing detection tools in several multimodal transcriptomic cancer datasets and provide recommendations based on those results. We found sample mispairs in three out of five datasets. Lastly, a high-grade serous ovarian carcinoma dataset is reconstructed: mispairs are resolved between several RNA-seq modalities and the connection between the transcriptomic data and patient metadata is reestablished by estimating the genetic ancestry from the transcriptomic data and comparing that with the metadata.

## Introduction

Sample mispairing caused by mislabeling or sample swaps is a documented problem in next-generation sequencing (NGS) studies. Lohr et al. [1] found that 40% of 45 transcriptomic datasets had sample mislabeling errors, and Grimm et al. [2] estimates an error rate of around 1% in NGS data. Resolving sample swaps has also been shown to improve the ability to detect biologically relevant patterns [3]. In severe cases, sample swaps can lead to incorrect conclusions and wasted resources, and therefore it is important to identify and correct sample swaps.

Many tools have been developed to identify sample swaps in genomic data, often with specific use-cases in mind such as single-modality longitudinal or spatial data, tumor vs germline, or whole-genome sequencing (WGS). Existing tools address sample-pairing errors in different settings, but their relative performance in multimodal cancer transcriptomic data remains unclear. A benchmark assessing tools for the detection of cross-contamination in NGS cancer data was published in 2024 [4], but there is no similar benchmark for the detection of sample mispairing.

This study evaluates sample mispairing identification tools in multimodal transcriptomic cancer datasets. Mispaired samples are samples that have been incorrectly attributed to the same donor. In paired NGS data, there is an opportunity to verify the sample pairing as part of the quality control (QC) process. Cancer data adds an extra challenge because tumor cells have greater genomic variability and gene expression variability than normal tissue cells from the same donor.

A review of the literature identified many tools to assess sample mispairs, swaps, or duplicates. reGenotyper [5] is built for high-throughput applications and needs many samples to work well. MBV [6] is built with expression quantitative trait loci (eQTL) detection in mind, comparing BAM (aligned reads) with VCF (variant calls). BIGRED [7] has the ability to detect swaps even when there is low coverage, but it relies on WGS. DoppelgangR [8] was developed to identify duplicate expression profiles in public databases. BAM-matcher [9] was included in this benchmark, but the authors were unable to get the software working. Next, BAMixChecker [10] ran after upgrading it to Python 3, however, no usable results were obtained since the program never predicts a matching pair of samples. The remaining tools are included in the benchmark results: Conpair [11], CrosscheckFingerprints [12], HYSYS [13], NGSCheckMate [14], ntsm [15],

OmicsPrint [16], Peddy [17], Somalier [18], TimeAttackGenComp [19] and Vireo [20]. Where possible, variant calls are used as input for the tools, and when necessary aligned reads (BAMixChecker, Conpair, Somalier) or raw reads (ntsm) are used as input instead.

There are three parts to this study: First, pseudobulk experiments with known sample pairings are used to benchmark sample mispairing identification tools in a controlled setting. This is needed because accuracy can only be evaluated when the ground truth of the correct sample pairs is known. Second, paired bulk and single-cell RNA-sequencing (RNA-seq) data from several datasets are used to evaluate the tools in a realistic setting. Lastly, we reconstruct a high-grade serous carcinoma (HGSOC) dataset with several mispairing issues: diagnosing and resolving mispairs from two bulk RNA-seq modalities, checking for sample swaps in pooled RNA-seq data, and aligning the RNA-seq sample ids with the metadata sample ids using ancestry inference.

The datasets used in this benchmark are listed in Tab. 1. The raw data and metadata can be found in the Sequence Read Archive (SRA) using the accession lists in the supplementary information. The pseudobulk experiments used all listed (non-pooled) single-cell and single-nucleus RNA-seq data.. The real data experiments tested the first four listed datasets and the dataset reconstruction is done on the HGSOC dataset listed last.

**Table 1.** The datasets and RNA-seq modalities used in this work, including the average read depth for each modality and the number of samples. The average read depth was calculated using Mosdepth [21]. The datasets above the dashed line were used in benchmarking and the last dataset is used as a case study. rRNA-stands for rRNA depleted and poly A+ for poly A enriched. ‘diss bulk’ is short for dissociated bulk, which is to say tissue was dissociated into individual cells and bulk sequenced as opposed to the traditional bulk sequencing, referred to as ‘chunk bulk’ here.

| **Dataset** | **Modality** | **Read depth** | **Sample size** |
|---|---|---|---|
| High-grade glioma | bulk | 33 | 18 |
| High-grade glioma | single-cell | 23 | 19 |
| HGSOC (pilot) | rRNA- chunk bulk | 24 | 8 |
| HGSOC (pilot) | rRNA- diss bulk | 22 | 8 |
| HGSOC (pilot) | poly A+ diss bulk | 32 | 8 |
| HGSOC (pilot) | single-cell | 566 | 8 |
| Low-grade glioma | bulk | 2 | 8 |
| Low-grade glioma | single-cell | 19 | 22 |
| Wilms' tumor | bulk | 111 | 42 |
| Wilms' tumor | single-nucleus | 326 | 40 |
| HGSOC (new) | rRNA- chunk bulk | 14 | 36 |
| HGSOC (new) | poly A+ diss bulk | 13 | 32 |
| HGSOC (new) | pooled single-cell | 1065 | 37 |

## Results

The preprocessing and benchmarking workflows are shown condensed in Fig. 1 and fully in Fig. S1. Sample mispair detection tools receive raw reads, aligned reads, or variant calls as input. Each tool calculates a distinct sample similarity metric for pairs of samples. Some tools report a sample similarity metric that can be thresholded to classify sample pairs as matches or mismatches. For others, an algorithm is applied to determine whether a particular pair of samples matches, does not match, or is inconclusive. Most tools compare genetic variants between two samples to determine their similarity, which also includes identity-by-state methods. Despite the apparent similarity, each tool has a distinct approach (e.g., focusing on homozygous markers, only looking at SNPs, or considering the variant allele fractions). Some methods take a completely different approach, looking at the linkage disequilibrium (CrosscheckFingerprints), and counting k-mers (ntsm).

**Fig 1.**
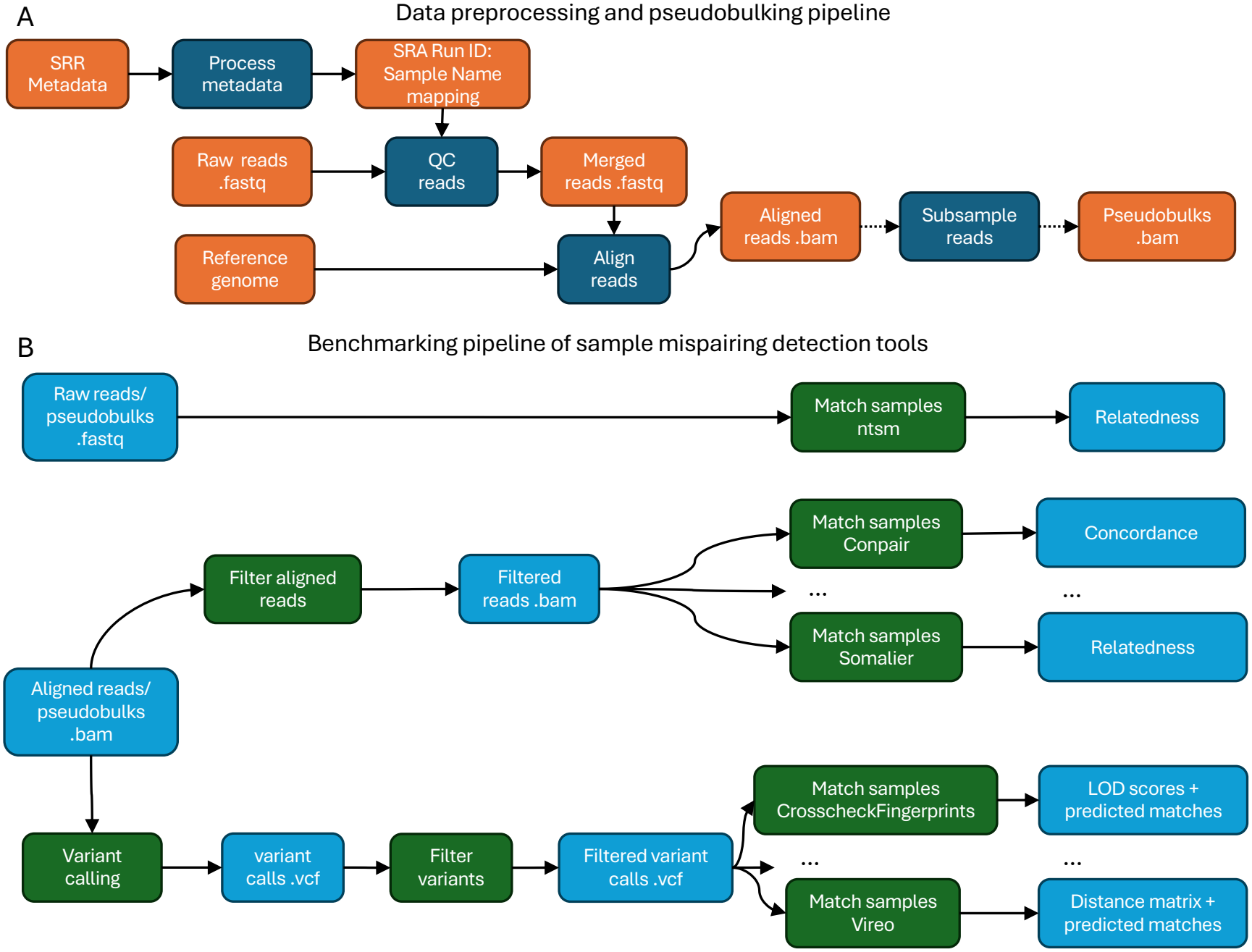
A) Workflow diagram of the data preprocessing and pseudobulking pipeline: samples are QC’ed with fastp, reads are aligned with STAR, and reads are sampled with subset-bam. Orange boxes represent data and blue boxes represent processes applied to those data. Solid arrows indicate mandatory steps and dashed arrows indicate optional steps. B) Workflow diagram of the benchmarking pipeline: Tools requiring raw reads as input are in a separate pipeline (top). Tools requiring aligned reads follow the middle branch: reads are filtered and then fed into the sample mispair detection tools. Tools accepting variant calls follow the bottom branch: variants are called and filtered and then fed into the various tools.

### Pseudobulk experiments with known sample pairings

The benchmarking results of sample similarity heatmaps of high-grade glioma pseudobulks are shown in Fig. 2A for each tool and for an increasing number of unique molecular identifiers (UMIs) that were included in the pseudobulks. Each heatmap compares two sets of pseudobulks drawn from the same set of single-cell RNA-seq samples. Unless otherwise stated, the samples in each sample similarity heatmap plot are ordered such that the ground truth sample matches lie on the diagonal. The patterns in the plots with fewer than 500k UMIs are somewhat masked by the shared heatmap scales for each row. The diagonal of correctly identified sample matches is clearly visible in most heatmaps with 500k UMIs, and seven of the tools research an F1 score of 1. Recall, precision, and F1 score are calculated by comparing the tools’ predictions of each sample pair (match, no match, or inconclusive) to the ground truth. Note that inconclusive calls are counted as ‘no match’ for the purpose of calculating the recall, precision, and F1 score. Performance generally improved as the number of UMIs increased.

**Fig 2.**
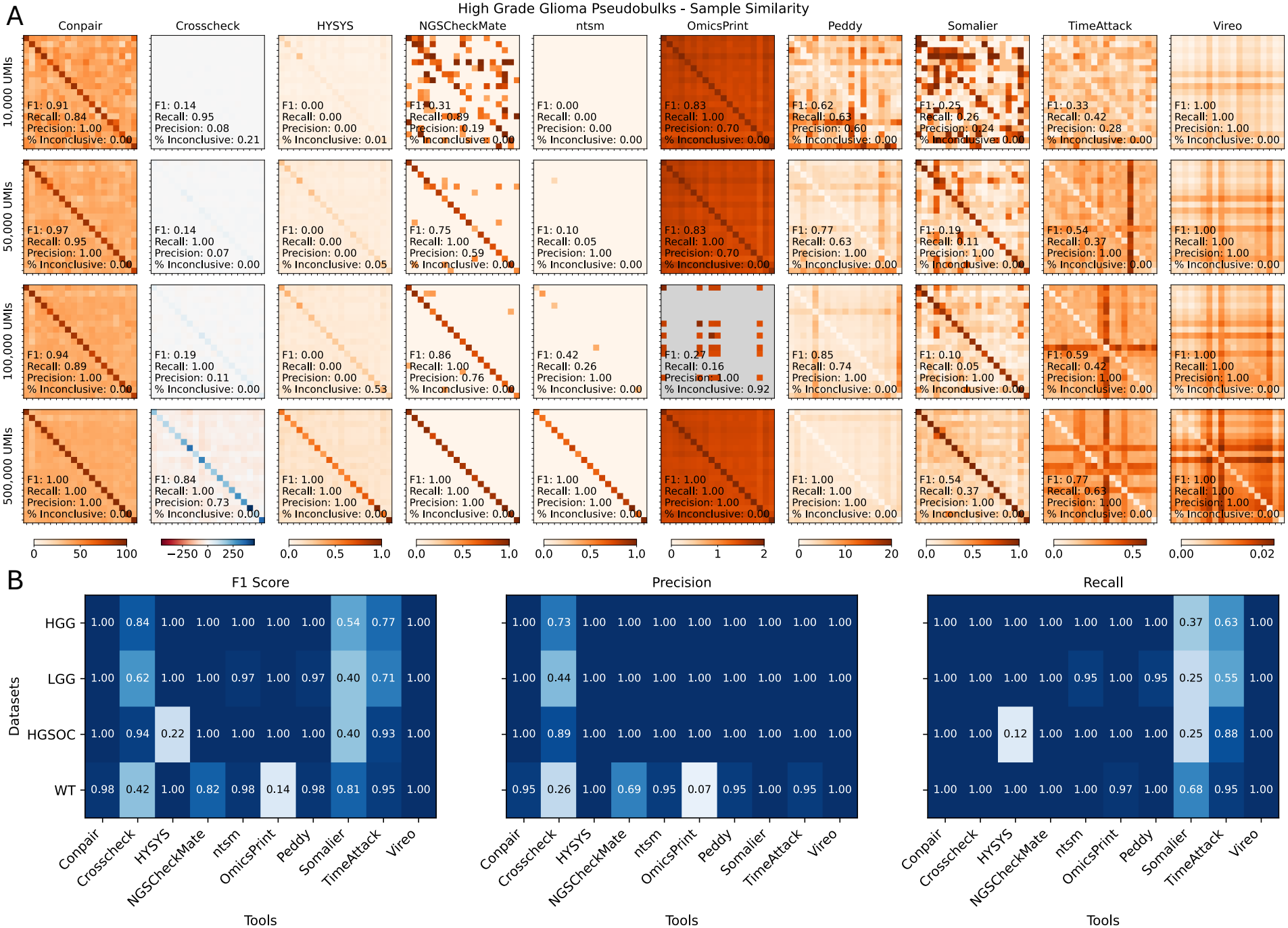
A) Sample similarity heatmaps comparing two sets of pseudobulks drawn from the same set of high-grade glioma single-cell RNA-seq data for the different tools and increasing number of UMIs in each pseudobulk. Grey indicates no sample similarity metric was calculated. The ground-truth sample matches lie on the diagonal. Performance metrics comparing the tools’ calls (match, no match, or inconclusive) to the ground truth are printed in the lower left corner of each heatmap. Any inconclusive calls were counted as “no match” for the calculation of the precision, recall, and F1 score. B) Performance metrics comparing the tools’ calls to the ground truth for the different tools and different datasets (High-grade serous ovarian cancer (HGSOC), high-grade glioma (HGG), low-grade glioma (LGG), Wilms’ tumor (WT)) with 500k UMIs in each pseudobulk.

Performance metrics for all tools and pseudobulk datasets are shown in 2B. The corresponding heatmaps can be found in the supplementary information, Fig. S2. Conpair, ntsm, Peddy, and Vireo stand out as achieving near perfect F1 scores for all pseudobulk datasets. Most tools achieved high precision or high recall, but did not consistently achieve both. For Conpair, OmicsPrint, Peddy, Somalier, and TimeAttackGenComp thresholds had to be set on the sample similarity above or below which to call a pair of samples matching, details in the Methods section.

One reason why tools might be doing worse on the low-grade glioma or high-grade glioma pseudobulks is that they have the lowest read depths. On the other hand, the Wilms’ tumor pseudobulks have great depth, but there is a large number of samples to compare. Finally, some tools like CrosscheckFingerprints might still improve with more data (more than 500k UMIs).

### Real data

We next evaluated the tools using real cancer datasets comprising different RNA-seq modalities. The sample similarity heatmaps comparing 19 single-cell and 18 bulk RNA-seq high-grade glioma samples from the same donors are shown in Fig. 3A. Again, the samples are ordered such that the expected matches lie on the diagonal and any mismatches should appear off the diagonal. For tools that used raw or aligned reads as input, the lower-bound read depth was not varied (Conpair, ntsm, Somalier). The different lower bound read depth cut-offs have a different effect on the heatmaps from the different tools; There is no single lower bound read depth that is ideal for all tools. The tools predict different sets of sample matches, but looking for consensus between the predictions, no sample mismatches are found.

**Fig 3.**
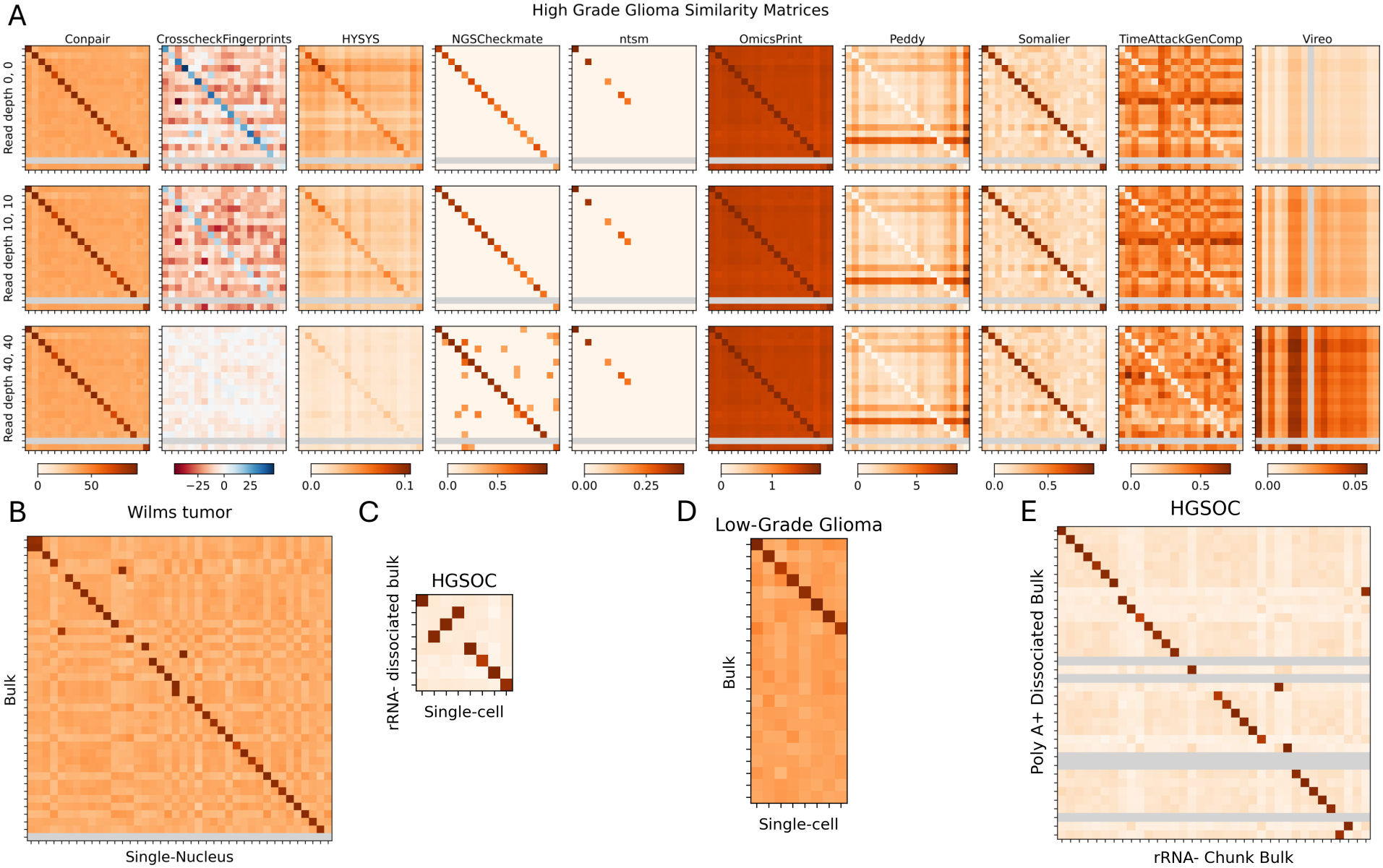
Sample similarity heatmaps comparing two different RNA-seq modalities: A) High-grade glioma single-cell (x-axis) compared to bulk RNA-seq (y-axis) samples from the same donors for different tools and lower-bound read depth cut-offs, B) Conpair heatmap of Wilms’ tumor single-nucleus compared to bulk RNA-seq, C) high-grade serous ovarian cancer (pilot) single-cell compared to rRNA-dissociated bulk RNA-seq, D) low-grade glioma single-cell compared to bulk RNA-seq. Figures B-E) do no have lower-bound read depth filters, E) high-grade serous ovarian cancer (new) rRNA-bulk chunk compared to polyA+ dissociated bulk RNA-seq. In all heatmaps samples are ordered such that the expected matches lie on the diagonal and any mismatches should appear off the diagonal. Grey bars indicate missing samples.

Conpair’s sample similarity heatmaps of Wilms’ tumor, HGSOC, and low-grade glioma are shown in Fig. 3B-D. The Wilms’ tumor and both HGSOC heatmaps clearly show instances of sample mispairing. The low-grade glioma dataset was the hardest dataset to assess since many tools struggled with the relatively low read depth of this dataset. Conpair had the clearest signal that showed no evidence of sample mismatches. Sample similarity heatmaps for all of these datasets and all of the tools can be found in the supplementary information, Fig. S3-Fig. S7. Conpair appeared to provide the highest signal-to-noise ratio across datasets, although its computational cost was substantially greater than that of the other tools. Many other tools yield similar predictions, though it can be hard to trust the predictions of a single tool. For this reason, the consensus between tools was used in this section to make the final decision on (mis)pairs.

### Restoration of a HGSOC dataset

The dataset under investigation was established to contain three RNA-seq modalities: rRNA-chunk bulk, poly A+ dissociated bulk, and pooled single-cell. The sample pairing of the two bulk RNA-seq modalities was assessed in Fig. 3E, and a large number of sample mispairs was found due to the propagation of an off-by-one error. Heatmaps for the other tools and the list of predicted sample matches can be found in Fig. S7 and the inferred sample matches can be found in S1 Table.

Since many tools are yielding similar predictions, the computational cost associated with running each tool is of interest. The computational costs broken down by Nextflow processes are shown in Tab. 2. The processes are grouped by input type (variant calls, aligned reads, raw reads), and the dashed lines separate the shared processes from the sample mispair detection tool processes. Conpair required 1,152 processes for the 32-by-36 sample comparison, whereas TimeAttackGenComp failed to complete on this dataset. For tools that use variant calls as input, genotyping is the most expensive step. An estimate of the total memory and time used per tool is added in Tab. 3. Conpair is computationally more expensive than the other tools by a wide margin. This is in large part due to a new process having to run for each pair of samples without caching of intermediate results that are used many times. In particular, there is an open pull request from 2025 in the Conpair Github repository to add parallel processing capabilities. Somalier is the cheapest tool in terms of RAM used and ntsm is the quickest. When variants are needed, running all of these tools and identifying consensus in the results may be justified.

**Table 2.** The computational burden of the Nextflow processes used to run the benchmarking workflow on the HGSOC dataset that is being restored. n counts the number of (potentially parallel) processes that were run, mem shows the memory that assigned to each process, %cpu shows the percentage of cpu that was used, realtime shows the time the process took to run, peak rss shows the peak memory usage. All of these quantities are averages over all the n processes. The total memory used was calculated by multiplying the number of processes n with the assigned memory and the %cpu used. The total time was calculated multiplying the number of processes n by the realtime. The top block lists all processes used for sample mismatch detection tools that accept variant calls as input. The middle block lists all processes for tools that need aligned reads as input. And, the bottom block lists all processes for tools that use raw reads as input. The dashed lines separate shared processes for each block from the processes that use them. Note that TimeAttackGenComp failed to run on this dataset.

| **Process name** | **n** | **mem<br>(GB)** | **%cpu** | **realtime<br>(s)** | **peak_rss<br>(GB)** | **total mem<br>(GB)** | **total time<br>(hr)** |
|---|---|---|---|---|---|---|---|
| PREPARE_INPUTS | 68 | 8 | 92.8 | 23.9 | 0.03 | 505 | 0.45 |
| SNP_REF | 1 | 8 | 70.1 | 41.6 | 0.7 | 5.6 | 0.01 |
| GENOTYPE_IND | 68 | 64 | 102 | 392 | 0.5 | 4,439 | 7.40 |
| MERGE_AND_FILTER | 1 | 8 | 56.5 | 571 | 0.6 | 4.5 | 0.16 |
| CROSSCHECK | 1 | 16 | 139 | 67.0 | 0.6 | 22.2 | 0.02 |
| HYSYS | 1 | 32 | 50.5 | 441 | 1.3 | 16.2 | 0.12 |
| NGSCHECKMATE | 1 | 8 | 8.6 | 222 | 0.2 | 0.69 | 0.06 |
| OMICSPRINT | 1 | 24 | 99.6 | 186 | 5.8 | 23.9 | 0.05 |
| PEDDY | 1 | 8 | 9.7 | 84.0 | 0.7 | 0.78 | 0.02 |
| TIMEATTACK | 0 | 32 | - | - | - | - | - |
| VIREO | 1 | 64 | 27.1 | 468 | 27.1 | 17.3 | 0.13 |
| FILTER_BAM | 68 | 8 | 95.7 | 313 | 0.02 | 521 | 5.91 |
| CONPAIR_PILEUP | 68 | 32 | 74.6 | 78.9 | 0.6 | 1,623 | 1.49 |
| CONPAIR_VERIFY | 1152 | 8 | 47.9 | 3.5 | 0.1 | 4,414 | 1.1 |
| SOMALIER_EXTRACT | 68 | 16 | 95.0 | 98.0 | 0.5 | 1,034 | 1.85 |
| SOMALIER_RELATE | 1 | 16 | 2.7 | 27.2 | 0.06 | 0.43 | 0.03 |
| NTSM_COUNT | 68 | 32 | 184 | 231 | 0.2 | 4,004 | 4.36 |
| NSTM_EVAL | 1 | 8 | 36.2 | 14.3 | 0.1 | 2.9 | 0.004 |

**Table 3.** Computational burden estimate per tool including all processes needed to run each tool. E.g. For CrosscheckFingerprints that combines the processes PREPARE INPUTS, SNP REF, GENOTYPE INDIVIDUAL, MERGE AND FILTER, AND CROSSCHECKFINGERPRINTS. TimeAttackGenComp failed to run on the HGSOC dataset.

| **Tool** | **total memory (GB)** | **total time (hr)** |
|---|---|---|
| CrosscheckFingerprints | 4,976 | 8.0 |
| HYSYS | 4,970 | 8.1 |
| NGSCheckMate | 4,955 | 8.1 |
| OmicsPrint | 4,978 | 8.1 |
| Peddy | 4,955 | 8.0 |
| TimeAttackGenComp | - | - |
| Vireo | 4,971 | 8.2 |
| Conpair | 6,558 | 8.5 |
| Somalier | 1,555 | 7.8 |
| ntsm | 4,007 | 4.4 |

### Genetic ancestry

Samples originating from the same donors have been identified successfully and the only missing piece is the connection to the metadata: which of the sets of RNA-seq modalities’ sample ids correspond to the sample ids in the metadata? The estimated genetic ancestry proportions for each modality and sample can be found in Fig. S8. PCA plots of those estimates are shown in Fig. 4B superimposed on genetic ancestry references from the 1000 Genomes Project [22]. The data from this study have a black edge. The location in the PCA plot shows the inferred ancestry estimate and the color shows the ancestry label according to the metadata. For poly A+ dissociated bulk there are nine donors whose inferred ancestry group does not match the ancestry information from the metadata, while for rRNA-chunk bulk all the donors’ inferred ancestry groups match the metadata. Hence, the rRNA-chunk bulk RNA-seq samples are concluded to be labeled correctly (matching the metadata), and the poly A+ dissociated bulk RNA-seq samples have been mislabeled. The inferred matching samples between the two RNA-seq modalities and the metadata are shown in S1 Table.

**Fig 4.**
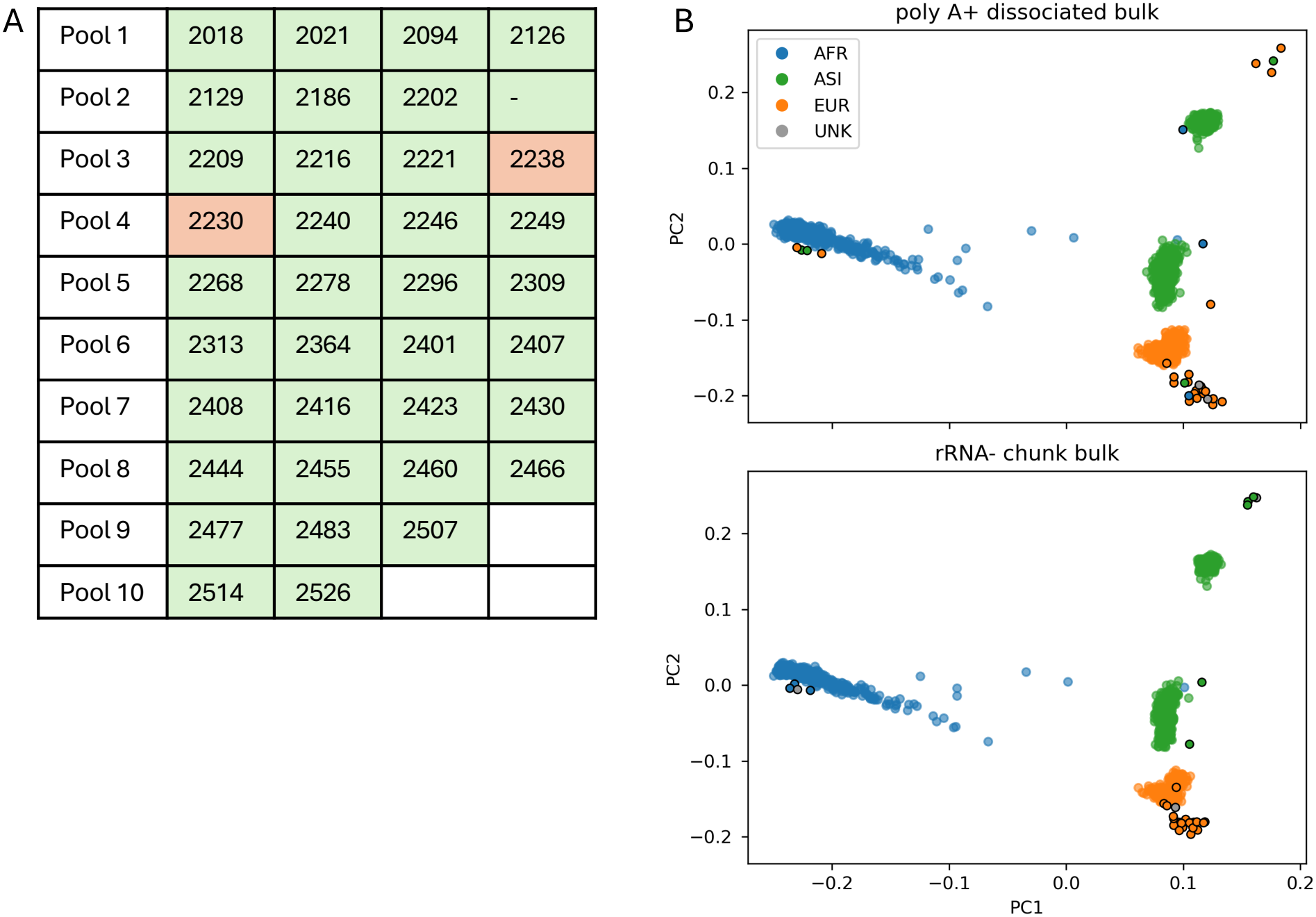
A) Demultiplexing of the pooled scRNA-seq data with rRNA-bulk RNA-seq as a reference without assuming which samples are in each pool reveals a sample swap between pools 3 and 4. B) PCA plot of 1000G references (transparent points) and HGSOC donors (opaque points with a black border) comparing the metadata ancestry labels (color) against the inferred ancestry labels (location in PCA space). AFR = African, ASI = Asian, EUR = European, UNK = unknown.

## Discussion

Twelve sample mispairing detection tools were benchmarked in both pseudobulk and real data. A summary of our benchmark conclusions is shown in Fig. 5. Two (BAMixChecker and BAM-matcher) were discarded due to performance issues. In pseudobulks, Conpair, NGSCheckMate, ntsm, Peddy, and Vireo turned out to be the most robust to sample size and different datasets. Performance was less consistent across tools and datasets in the real-data experiments: ntsm, OmicsPrint, and TimeAttackGenComp fail to perform on specific datasets. However, none of the tools failed for any specific modality (bulk, single-cell, or single-nucleus RNA-seq). Since we do not know the ground-truth sample matches in real datasets, we cannot say with certainty which tools are the most accurate. In the reconstructed HGSOC dataset, we have an additional validation step through the genetic ancestry inference, and we used that to substantiate Conpair’s predicted sample matches as true matches. We then compared those matches with the other tools’ predictions to assess their performance; see Fig. 5.

**Fig 5.**
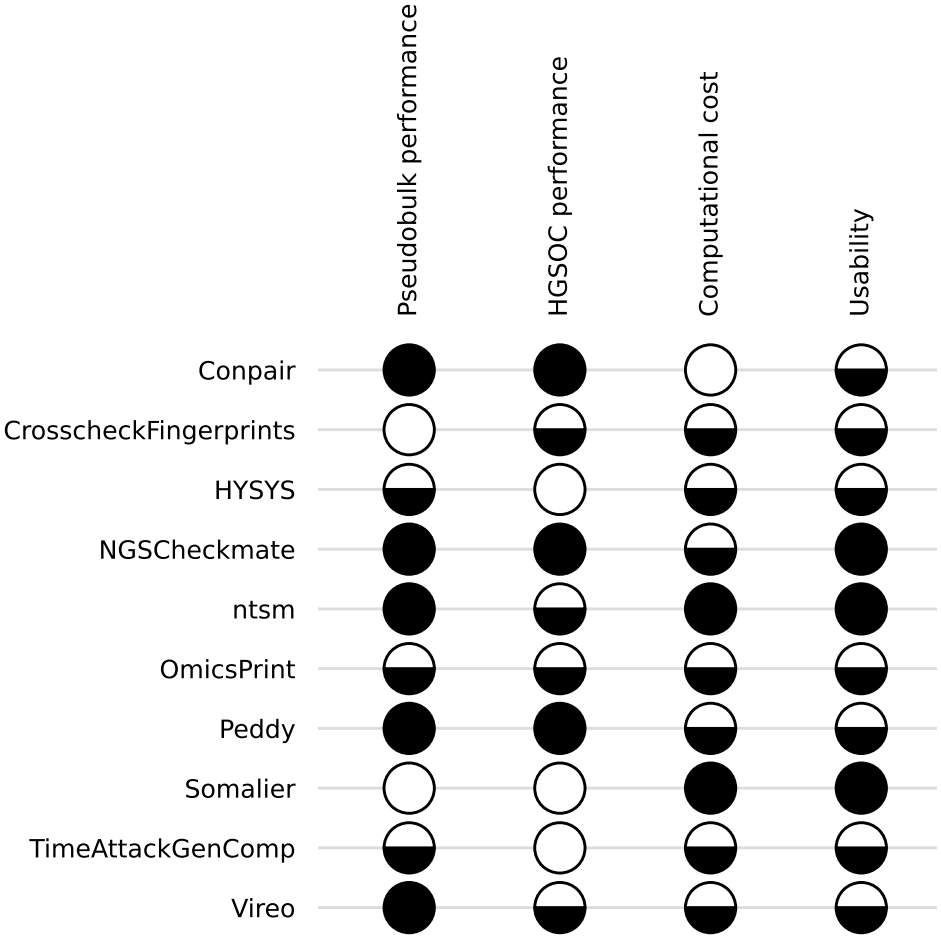
Benchmark summary by methods (rows) and evaluation criteria (columns). A filled circle indicates the method was superior, a half-filled circle indicates the method was satisfactory, and an empty circle denotes the method was unsatisfactory. A detailed rubric is provided in the Methods section.

Although Vireo performed well in the pseudobulk experiments, the less distinct patterns in its real-data heatmaps complicate the interpretation of its predicted sample matches. Vireo’s approach that makes it give good predictions despite a noisy heatmap is the linear sum assignment algorithm; It assumes that only one sample from each modality can match each other, and this is enforced by minimizing the diagonal of the heatmap matrix by permuting rows and columns. This assignment strategy may be applicable to other similarity matrices in the benchmark, provided that the underlying one-to-one matching assumption is satisfied.

In the low-grade glioma dataset, which had the lowest read depth, Conpair, Peddy, and Somalier produced the clearest sample-matching results. In the Wilms’ tumor dataset, which has the largest number of samples and compares single-nucleus with bulk RNA-seq data, Conpair, HYSYS, and Somalier performed well. In three out of five tested datasets (HGSOC (pilot), HGSOC (new), Wilms’ tumor) sample mispairs were found, and two (low-grade glioma and high-grade glioma) were found to contain no sample mispairs. The reconstructed HGSOC dataset contained numerous sample swaps, which we resolved using the benchmarking pipeline. And finally, genetic ancestry inference was used to align the sample IDs from the RNA-seq data with the metadata. The HGSOC dataset reconstruction was successful: all RNA-seq modalities and metadata are now aligned on sample ID.

For most tools, the variant calling part of the mispair detection process is the most costly in terms of computational power and time. For analyses that already require variant calling, we recommend evaluating agreement among the predictions of the seven tools implemented in our Nextflow pipeline. Alternatively, with aligned reads, Conpair and Somalier are both good options depending on compute resources. For datasets with sufficient read depth that do not otherwise require variant calls or aligned reads, ntsm may be a practical option.

## Methods

### Experimental methods

The experimental methods were based on those of a previous pilot study [23]. Briefly, samples were collected from 38 patients with HGSOC and minced into 1 mm pieces. Samples from each patient were sequenced in three different ways: (1) tumor chunks were depleted of ribosomal RNA and bulk sequenced, (2) dissociated samples were poly-A enriched and bulk sequenced, and (3) dissociated samples were pooled together into ten batches of up to four samples and single-cell sequenced. All bulk samples were sequenced on an Illumina NovaSeq 6000 system using S1 Reagent Kits v1.5 (200 cycles) and S2 Reagent Kits v1.5 (200 cycles).

Multiplexed single-cell samples were prepared for sequencing using the 10X Genomics 3’ CellPlex Kit and sequenced on an Illumina NovaSeq 6000 system using an S2 Reagent Kit v1.5 (100 cycles).

### HGSOC data identification

The dataset reconstruction began with four directories of RNA-seq data lacking complete metadata. Some directories contained partial annotations describing their contents. Differential expression analysis (DEA) was used on the four folders to confirm our hypotheses about the RNA-seq modalities in each of the folders. Briefly, sequence alignment was performed using STAR, count matrices were constructed for each folder of samples, and DEA was applied using PyDESeq2. All of the folders were compared against each other. Two sets of two folders with very few differentially expressed genes between each other were found, suggesting that they came from the same modalities. Between the two sets, significant differences in expression levels in mitochondrial and histone genes and, to a lesser extent, in hemoglobin- and adipose-related genes were found. Therefore, one set was identified as rRNA-bulk RNA-seq data from tissue chunks, and one as poly-A+ bulk RNA-seq data from dissociated tissue.

In addition to aforementioned analysis, the differential expression results were compared to DEA from another HGSOC dataset containing the same RNA-seq modalities.

### Preprocessing and pseudobulking

The preprocessing steps to prepare the data for benchmarking are implemented in Nextflow. The workflow is shown in Fig. 1A. To start, the mapping between SRA Run IDs and sample names is extracted from the SRA run metadata. The quality of the raw reads for each sample is checked using fast [24]. The reads are aligned to a reference genome using STAR [25], and the data is ready for the sample mispairing detection pipeline. Additionally, to create pseudobulks at the read level, there is the option to subsample the aligned reads using subset-bam.

### Benchmarking

The benchmarking pipeline was implemented in Nextflow and takes as input aligned reads in BAM format and a reference genome (the same one that was used for sequence alignment). A condensed version of the pipeline workflow is shown in Fig. 1B and version true to the Nextflow pipeline architecture is shown in the supplementary information. There are two main paths in the benchmarking workflow. For any tools that accept variant calls as input, first variant calling and filtering (by quality score and read depth) is done using bcftools [26] and is limited to single-nucleotide polymorphisms (SNPs) at specific locations specified by a list from The 1000 Genomes Project [22]. The filtered variants are run through each of the sample mispair detection tools (CrosscheckFingerprints, HYSYS, NGSCheckMate, OmicsPrint, Peddy, TimeAttackGenComp, and Vireo), yielding unique sample similarity estimates of each tool and predicted sample matches. For tools that do not accept variant calls as input, aligned reads are filtered by quality score using samtools [26], and then passed onto sample mispair detection tools (BAMixChecker, BAM-matcher, Conpair, and Somalier). Lastly, the tool ntsm requires raw reads as input, and that tool was benchmarked outside of this pipeline. The previously generated pseudobulks were transformed from bam into fastq files using samtools for the ntsm pseudobulk experiments.

Some tools require a threshold to be set on the sample similarity in order to call samples matching or not. The chosen thresholds were: 80 for Conpair, 1.7 or OmicsPrint, 0 for Peddy, 1 for Somalier, and 0.05 for TimeAttackGenComp. The values were obtained by studying the original papers and are not finetuned for the specific datasets used here. All results are visualized and analyzed in Python. The performance of the tools in pseudobulks was quantitatively evaluated using the fraction of inconclusive calls, precision, recall, and the F1 score. The performance of the tools in real data was assessed more qualitatively by looking at the signal-to-noise ratio of the heatmaps and comparing the predictions among the different tools. Sample (mis)pair conclusions for each dataset are based on all tools.

### Demultiplexing

The genetic demultiplexing protocol from [27] was adapted to eliminate the assumption that we know which samples are present in each pool. This was done by first variant calling without references using cellSNP-lite [28] and then demultiplexing with Vireo [20] using all poly A+ dissociated bulk or all rRNA-chunk bulk as references and letting the software infer which samples are in each pool.

### Genetic ancestry inference

The methods presented by Johnson et al. [29] were used to estimate the genetic ancestry of the different RNA-sequencing modalities. The SNPs that were generated in the variant calling step of the benchmarking pipeline were compared with references from The 1000 Genomes Project [22]. The estimated ancestry proportions for each sample can be found in Fig. S8. The most represented ancestry group from the estimated ancestry proportions of each sample was used to compare the inferred ancestry with the metadata. We grouped East Asian and South Asian ancestry together to match the groups present in our metadata. Note that we are not identifying individuals using this method, but rather seeing which set of sample ID labels fits the data best. Genetic ancestry estimation has been used in the process of detecting sample contamination [30] before, although the information was used to control for differences due to ancestry in that case.

### Benchmark evaluation criteria

#### Pseudobulk performance

Average F1 score across pseudobulk datasets with 500k UMIs

- Superior: greater than 0.95
- Satisfactory: 0.75-0.95
- Unsatisfactory: less than 0.75

#### HGSOC performance

F1 score on the HGSOC restoration dataset of tool predictions compared to Conpair-inferred reference assignments (corroborated by the ancestry inference).

- Superior: greater than 0.95
- Satisfactory: 0.75-0.95
- Unsatisfactory: less than 0.75

#### Computational cost

- Superior: Top tool in computational time and CPU usage
- Satisfactory: Middle of the pack tool in computation time and CPU usage
- Unsatisfactory: Bottom tool in computation time and CPU usage

#### Usability

- Superior: The tool is available and well documented, and no additional work is needed to get the tool working.
- Satisfactory: Tool is available, reasonably well documented and requires some additional work to get working.
- Unsatisfactory: Tool is unavailable or not working.

## Code availability

The code for the identification of HGSOC data is available at https://github.com/greenelab/hgsoc_data_detective. The code for the processing of the raw

RNA-seq data, the creation of pseudobulks, and the comparison of the different sample-swap identification tools is available at https://github.com/greenelab/hgsoc-reconstruction. The code for genetic ancestry inference is available at https://github.com/greenelab/hgsoc-concordance-rotation.

## Acknowledgments

The authors thank Jay Gertz and Noel Kitchen (University of Utah) for their advice on the experimental design. The authors thank Adriana Ivich and Natalie Davidson (University of Colorado Anschutz) for their efforts to identify the data types.

## Supporting information

**S1 Complete benchmarking pipeline diagram**

**S2 Pseudobulk sample similarity heatmaps** Sample similarity heatmaps accompanying Fig. 2B for all pseudobulk datasets.

**S3 Wilms’ tumor sample similarity heatmaps** Sample similarity heatmaps for all benchmarking tools for the Wilms’ tumor dataset.

**S4 Low-grade glioma sample similarity heatmaps** Sample similarity heatmaps for all benchmarking tools for the low-grade glioma dataset.

**S5 HGSOC (pilot) sample similarity heatmaps** Sample similarity heatmaps for all benchmarking tools for the pilot HGSOC dataset.

**S6 Multimodal HGSOC sample similarity heatmaps** Sample similarity heatmaps from HYSYS for the pilot HGSOC dataset comparing all the different RNA-seq modalities.

**S7 HGSOC (new) sample similarity heatmaps** Sample similarity heatmaps for all benchmarking tools for the new HGSOC dataset.

**S8 Ancestry proportions barplot** Estimated ancestry proportions from ancestry inference of the new HGSOC datasets for both bulk RNA-seq modalities.

**S1 Table HGSOC sample matches** Matched sample IDs for the bulk RNA-seq modalities and the metadata for the new HGSOC dataset.

