## Supplementary Information for "Benchmarking sample mispairing detection tools in complex cancer datasets"

### Supplementary Information: Benchmarking sample mislabeling detection tools in complex cancer datasets

#### SI Nextflow pipeline

Figure S1 shows all parts of the benchmarking pipeline implemented in Nextflow. There are three Nextflow pipelines in total: one for the data preprocessing and pseudobulking, one for testing ntsm (since it requires raw reads as input), and one benchmarking all the other tools.

#### SII Pseudobulk results

Figure S2 shows the sample similarity heatmaps corresponding to the performance metrics in Fig. 2B of the main text. Notably, a good signal-to-noise ratio of the heatmap does not necessarily correspond to a good overall performance.

#### SIII Real data results

##### SIII.1 Wilms' tumor

Sample similarity heatmaps of Wilms' tumor comparing bulk to single-nucleus RNA-seq for all tested tools is shown in Fig. S3. Conpair, CrosscheckFingerprints, HYSYS, OmicsPrint, and Peddy show very similar patterns; all predicting a couple of mismatches as we see three clear off-diagonal squares lighting up. Note that any blue square denotes a match for Crosscheck and red denotes non-match.

##### SIII.2 Low-grade glioma

Sample similarity heatmaps of low-grade glioma comparing bulk to single-cell RNA-seq for all tested tools is shown in Fig. S4. Conpair, Peddy, and Somalier show the most convincing diagonal in their heatmaps, while CrosscheckFingerprints, HYSYS, and TimeAttackGenComp are a bit noisier, and NGSCheckMate, ntsm, and Vireo look unusable. Vireo only returns sample similarity scores for matching samples, hence the smaller heatmap. The low-grade glioma bulk data has the lowest average read depth of all the data considered in this study, which might contribute to the noisier results. Yet, we are confident there are no samples mismatches in this dataset.

##### SIII.3 HGSOC

Sample similarity heatmaps of the HGSOC pilot dataset comparing rRNA-depleted dissociated bulk to single-cell RNA-seq for all tested tools is shown in Fig. S5. All tools except for NGSCheckMate

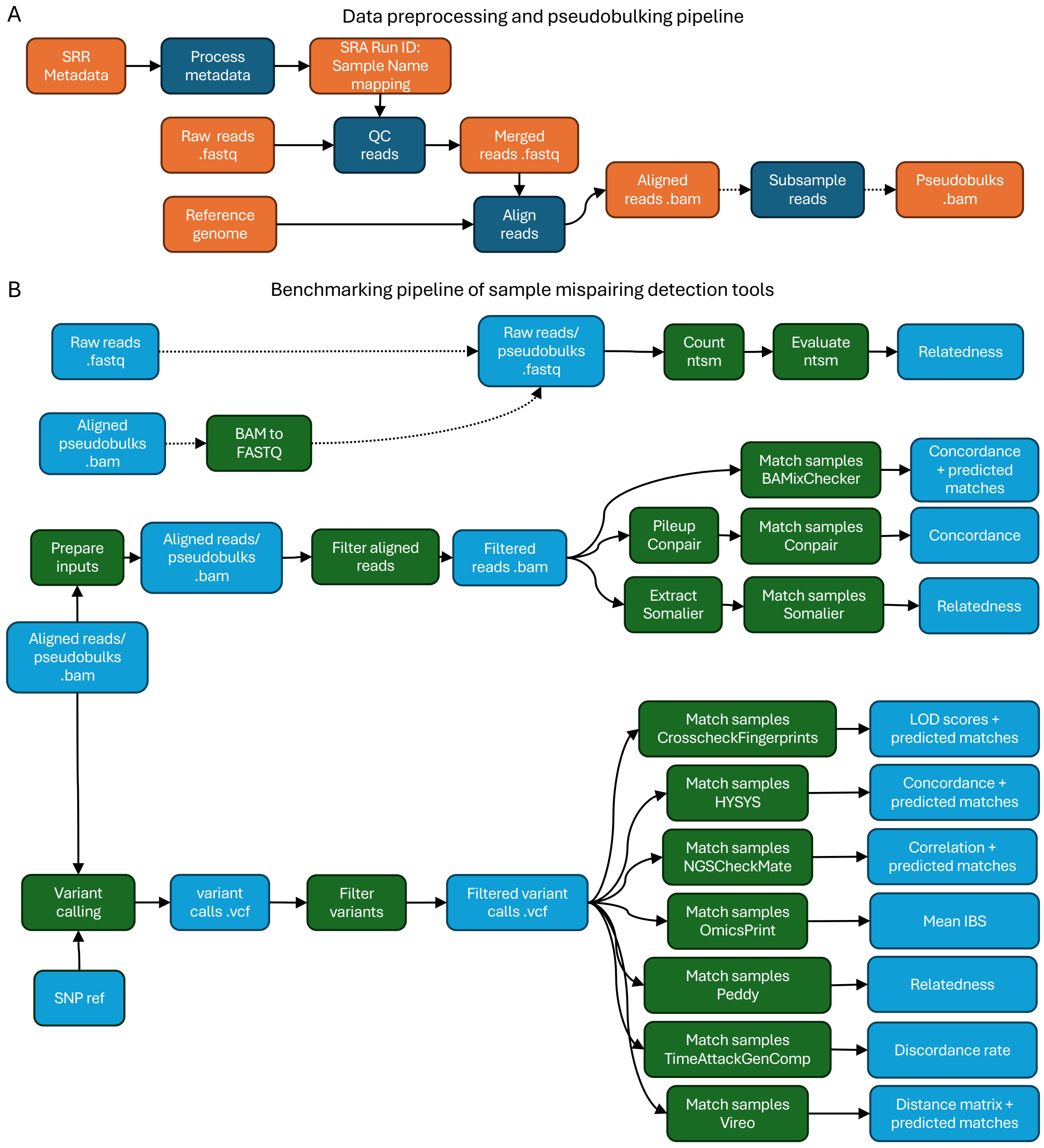


Figure S1: Complete overview of the benchmarking pipeline as implemented in Nextflow. A) Data preprocessing and pseudobulking pipeline. B) Benchmarking pipeline(s): The top pipeline tests ntsm, which requires raw reads as inputs. The bottom pipeline branches into two where one tests tools that accept variant calls (VCF) as inputs and the other tests tools that accept aligned reads (BAM) as inputs. The variant calls branch performs variant calling on aligned reads, filters those variants by quality score and read depth, and then feeds them into the tools to be tested (CrosscheckFingerprints, HYSYS, NGSCheckMate, OmicsPrint, Peddy, TimeAttackGenComp, and Vireo). The aligned reads branch filters the reads by quality score and then feeds them into the tools to be tested (BAMixChecker, Conpair, and Somalier).

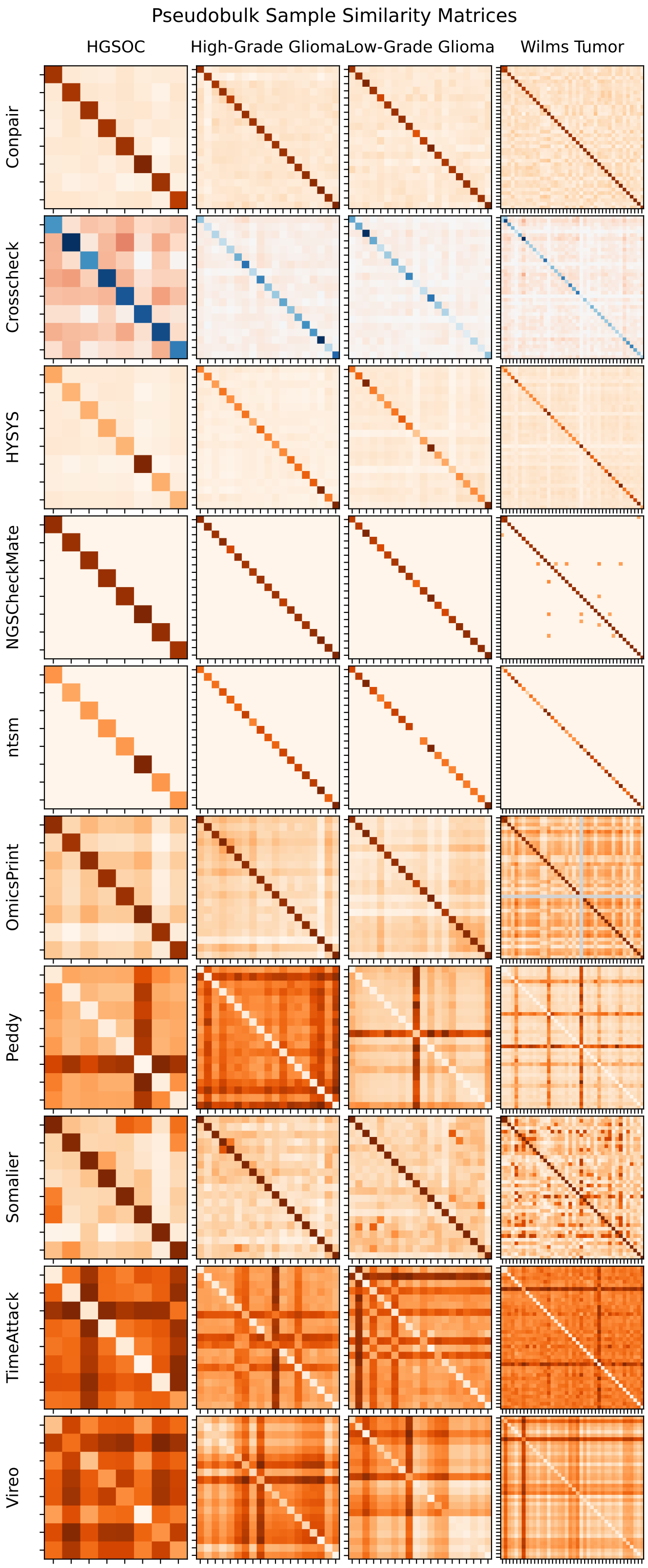


Figure S2: Sample similarity heatmaps for all datasets and tools in Fig. 2B of the main text, 500k UMIs and no lower bound read depth filter.

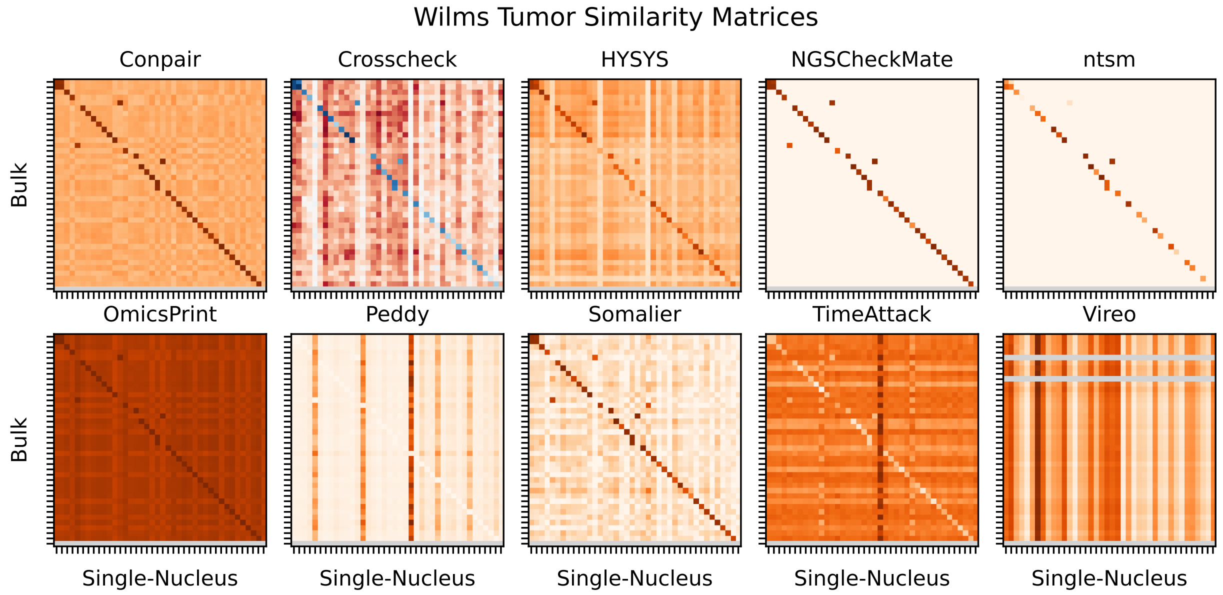


Figure S3: Wilms' tumor sample similarity heatmaps comparing bulk to single-nucleus RNA-seq for all tools.

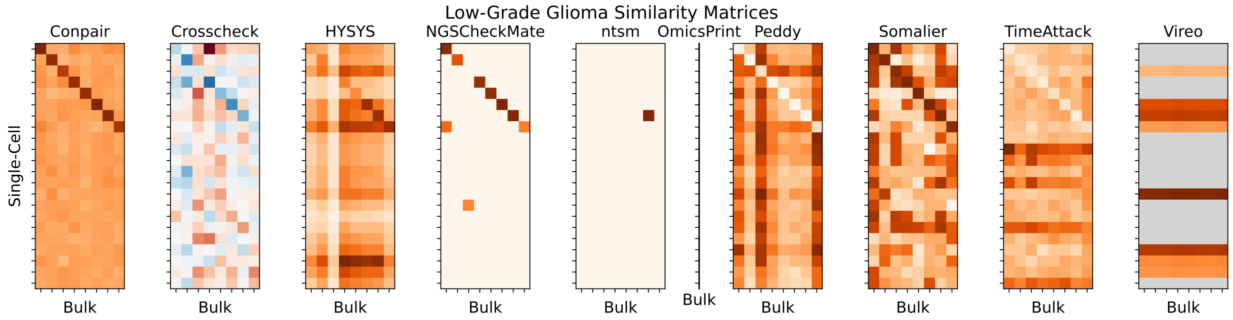


Figure S4: Low-grade glioma sample similarity heatmaps comparing bulk to single-cell RNA-seq for all tools.

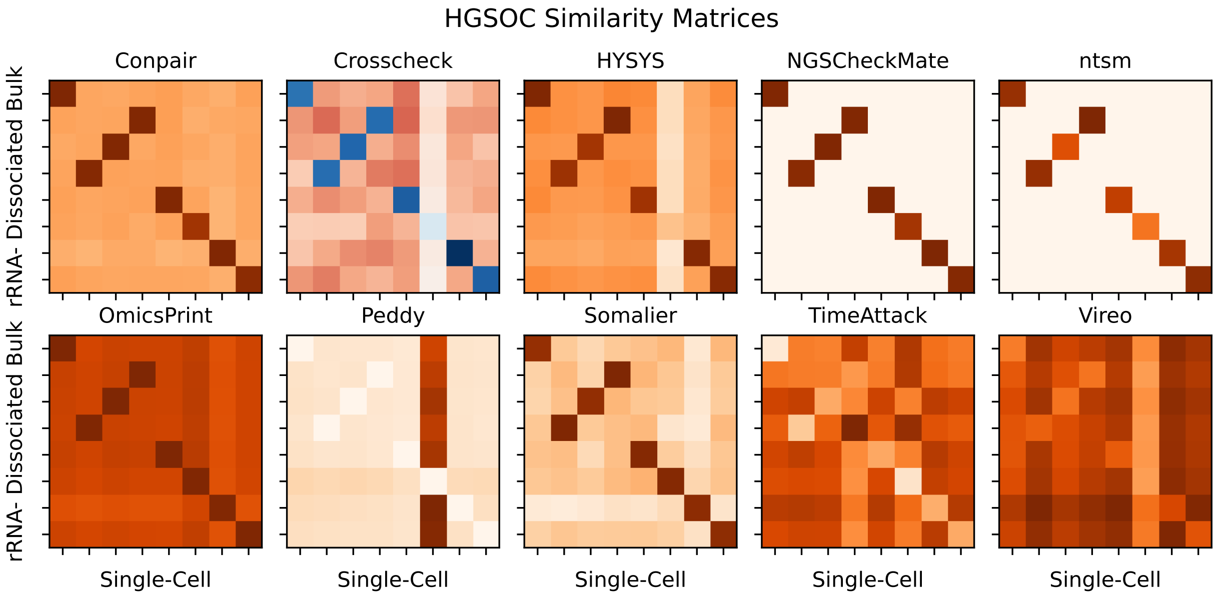


Figure S5: HGSOC sample similarity heatmaps comparing bulk dissociated rRNA- to single-cell RNA-seq for all tools.

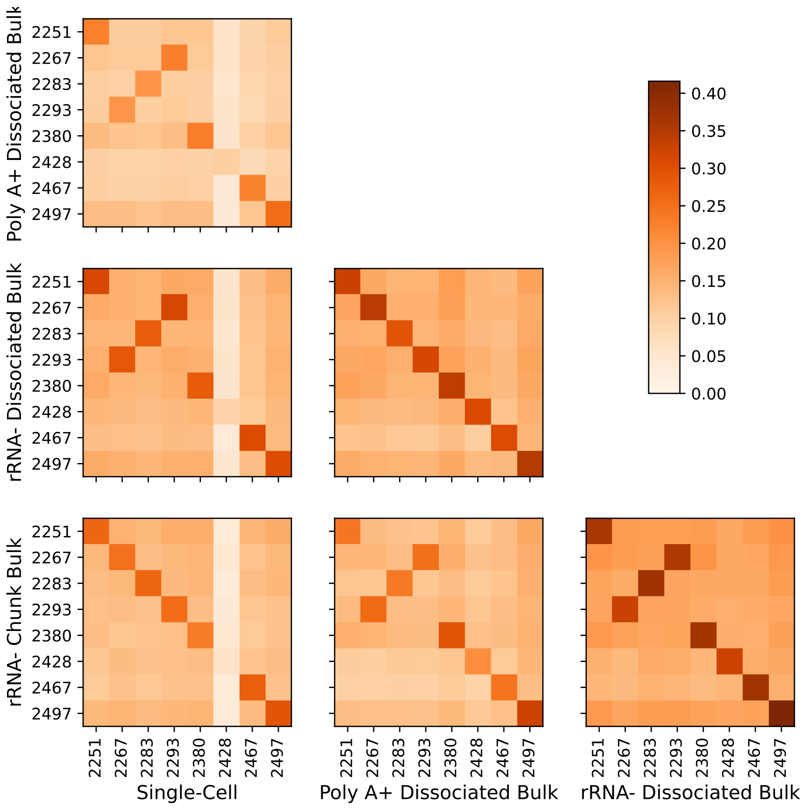


Figure S6: HYSYS sample similarity heatmaps for the HGSOC dataset comparing all the different modalities.

show the same sample-swap pattern albeit with varying levels of noise. All of the different RNA-seq modalities from this dataset are compared in Fig. S6 with the HYSYS heatmaps. The comparison of rRNA-depleted chunk bulk and single-cell samples agrees with the expected matches, while the dissociated bulk modalities show a sample was swapped with respect to the rRNA-depleted chunk bulk and single-cell modalities.

### SIV Dataset reconstruction

#### SIV.1 Sample mispairing

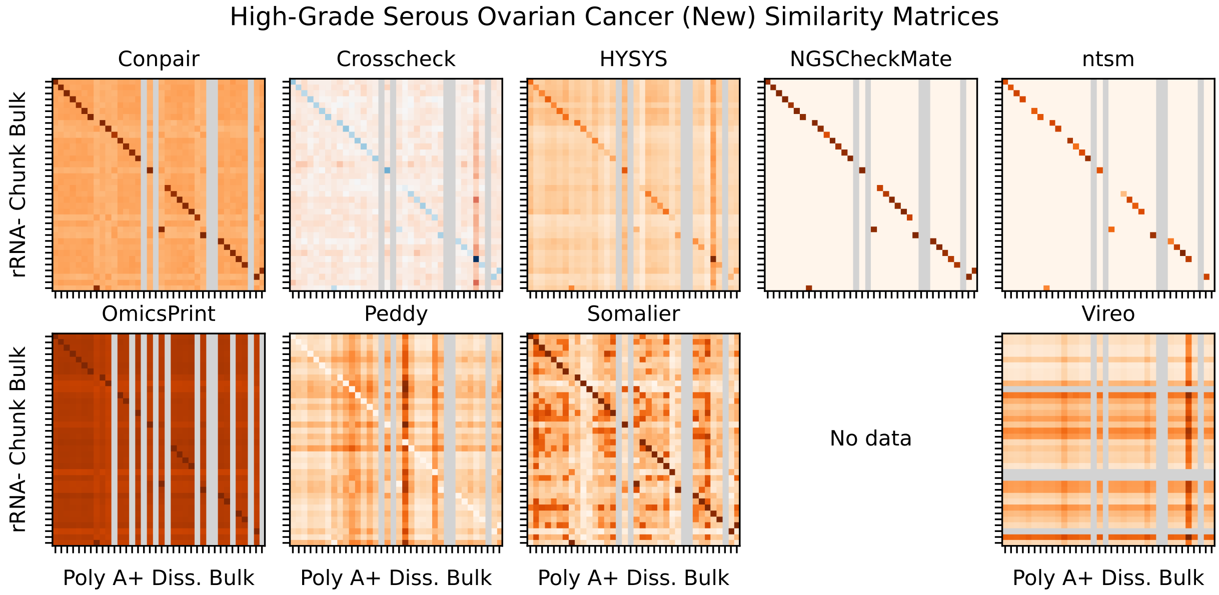


Figure S7: HGSOC sample similarity heatmaps comparing poly-A enriched dissociated bulk to rRNA-depleted chunk bulk for all the tested tools.

The sample similarity heatmaps for all the different tools applied to the HGSOC dataset are shown in Fig. S7. Conpair, CrosscheckFingerprints, HYSYS, OmicsPrint, Peddy, and Somalier heatmaps show very similar patterns, building confidence in the sample matches. TimeAttackGenComp failed to run, ntsm returns no matches, and NGSCheckMate and Vireo look more noisy. The predicted sample matches between the two bulk RNA-seq modalities are shown in Tab. S1.

#### SIV.2 Ancestry inference

The ancestry inference pipeline yields estimates of ancestry proportions that are shown in Fig. S8 for both bulk RNA-seq modalities. Note that it is not used to identify individuals, but rather to assess which set of sample ID labels (from rRNA- chunk bulk or poly A+ dissociated bulk) matches the metadata sample ID labels best. The inferred ancestry of each donor is taken as the most representative population in the ancestry proportions. As shown in the main text, Fig.4B, the rRNA- chunk bulk labels match the data well, whereas the poly A+ dissociated bulk labels do a worse job. The resulting matched sample IDs between the two bulk RNA-seq modalities and the metadata is shown in Tab. S1.

| Metadata sample ID | rRNA- chunk bulk sample ID | Poly A+ diss bulk sample ID |
|---|---|---|
| 2018 | 2018 | 2018 |
| 2023 | 2023 | 2023 |
| 2094 | 2094 | 2094 |
| 2126 | 2126 | 2126 |
| 2129 | 2129 | 2129 |
| 2186 | 2186 | 2186 |
| 2202 | 2202 | 2202 |
| 2205 | - | - |
| 2209 | 2209 | 2216 |
| 2216 | 2216 | 2221 |
| 2221 | 2221 | 2230 |
| 2230 | 2230 | 2238 |
| 2238 | 2238 | 2240 |
| 2240 | 2240 | 2246 |
| 2246 | 2246 | 2249 |
| 2249 | 2249 | - |
| 2268 | 2268 | 2278 |
| 2278 | 2278 | - |
| 2296 | 2296 | - |
| 2309 | 2309 | 2313 |
| 2313 | 2313 | 2364 |
| 2364 | 2364 | 2401 |
| 2401 | 2401 | 2407 |
| 2407 | 2407 | 2408 |
| 2408 | 2408 | 2416 |
| 2416 | 2416 | - |
| 2423 | 2423 | 2309 |
| 2430 | 2430 | 2423 |
| 2444 | 2444 | 2455 |
| 2455 | 2455 | 2460 |
| 2460 | 2460 | 2466 |
| 2466 | 2466 | 2477 |
| 2477 | 2477 | 2483 |
| 2483 | 2483 | 2526 |
| 2507 | 2507 | 2514 |
| 2514 | 2514 | 2507 |
| 2526 | 2526 | 2209 |

Table S1: Sample matches for the HGSOC dataset across metadata and both bulk RNA-seq modalities using the original sample IDs. Green rows were originally correctly labeled. Orange rows were originally incorrectly labeled. White rows have at least one missing sample.

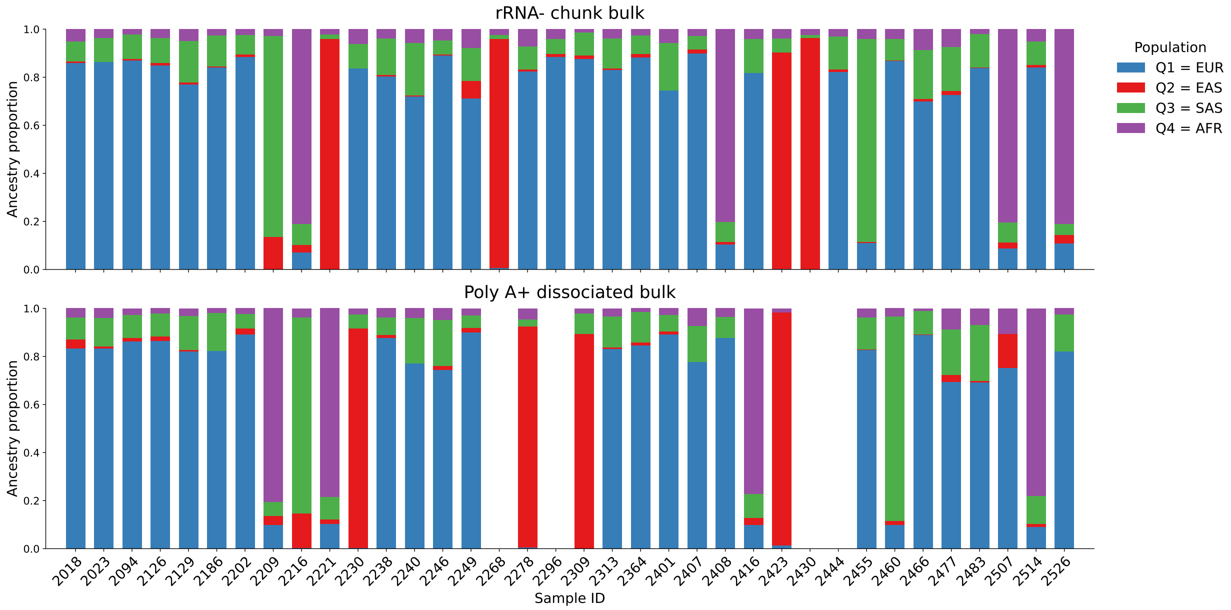


Figure S8: Ancestry proportions for all samples in both the rRNA- chunk bulk and poly A+ dissociated bulk modalities with the original sample labels.
